# Buprenorphine Restricts the Conformational Landscape of the *µ*-Opioid Receptor

**DOI:** 10.64898/2026.06.12.731880

**Authors:** Simone Aureli, Maria Bzówka, Nicola Piasentin, Massimo Fuss, Francesco Luigi Gervasio

**Affiliations:** School of Pharmaceutical Sciences, University of Geneva, Rue Michel-Servet 1, CH-1206 Geneva, CH; Institute of Pharmaceutical Sciences of Western Switzerland, University of Geneva, CH-1206, Geneva, CH; Swiss Bioinformatics Institute, University of Geneva, CH-1206, Geneva, CH; Department of Chemistry, University College London, London, WC1E 6BT, United Kingdom

## Abstract

Biased signaling and partial agonism in GPCRs are considered promising strategies to develop safer and more effective drugs. Various mechanisms have been proposed to explain these phenomena, including the ability of biased and partial ligands to promote specific receptor conformations able to engage distinct intracellular partners. Recent cryo-electron microscopy structures of the *µ*-opioid receptor (MOR) bound to both G-proteins and *β*-arrestins have shown clear structural differences shedding light into the putative signaling states. However, the accessibility of such conformations in the absence of the intracellular partner and the mechanisms by which ligands modulate their populations remain poorly understood. In the present manuscript, we employ extensive enhanced-sampling simulations to explore the free-energy landscapes of MOR bound to a biased partial agonist (buprenorphine) and a full unbiased agonist (endomorphin-1) showing that the ligands indeed stabilize distinct ensembles of active-like states. In particular, we observe that endomorphin-1, the natural ligand, explores two different ensembles of active-like conformations, including states compatible with either G-protein or *β*-arrestin1 recruitment, whereas buprenorphine restricts access to these conformations and favors a narrower subset of active states more prone to G-protein binding. Moreover, we analyze the allosteric pathways connecting the ligands to the different active-like states. Together, our findings provide a thermodynamic framework for understanding biased signaling at MOR and confirm that in MOR ligand efficacy and signaling preference emerge from the selective redistribution of receptor conformational populations even in the absence of the effector proteins. This perspective may guide the rational design of opioid therapeutics with improved signaling profiles and reduced adverse effects.

## Introduction

G protein-coupled receptors (GPCRs) are membrane proteins that belong to one of the largest and most pharmacologically relevant families of therapeutic targets, with more than 30% of all drugs acting on members of this receptor family^1–6^. GPCRs are known to populate different pre-active and active-like states^7,8^. Indeed, upon agonist binding, these receptors initiate intracellular signaling through interactions with multiple effector proteins, including various hetero-trimeric *G-proteins*^9^, *G protein–coupled receptor kinases* (GRKs also known as *β*ARK)^10,11^, and *β-arrestins*^12,13^. *Biased signaling* is the phenomenon where a ligand preferentially or exclusively activates one specific intracellular signaling pathway.^14–19^ Despite its promise for the development of safer and more effective drugs, the molecular mechanisms of *biased signaling* remain incompletely understood and actively debated. One of the most widespread explanation of this behaviour is the “conformational stabilization hypothesis”, where biased ligands induce and stabilize distinct receptor conformations. This is achieved through specific displacement patterns of transmembrane helices, particularly TM5, TM6, and TM7, and the dynamic repositioning of crucial structural elements like the intracellular helix 8 (H8). These dynamic structural rearrangements remodel the intracellular effector-binding cavity, sterically favoring the recruitment of one specific effector over another.^7,16,20–25^ The “phosphorylation barcode hypothesis” suggests instead that biased ligands expose different serine or threonine residues on the receptor’s intracellular loops or C-terminal tail, leading to selective recruitment of specific GRKs generating unique phosphorylation patterns that engage *β*-arrestins^20,26^. The “biased system” hypothesis suggests that biased signaling may result from differences in the expression of effector proteins. Notably, a ligand may cause an increased expression of specific GRKs or *β*-arrestin isoforms, which can steer signaling toward the *β*-arrestin pathway. On the other hand, insufficient levels of GRKs or *β*-arrestins may promote the recruitment of G-proteins instead.^7^. Finally, GPCRs can also form homodimers, heterodimers, or higher-order oligomers that inherently alter the receptor’s structural dynamics and allosteric communication, and can shift the signaling bias, for instance, by constitutively recruiting *β*-arrestins.^21,27,28^

The functional selectivity of the *µ*-opioid receptor (mOR) has been extensively studied in the global quest to develop safer analgesics and combat the ongoing opioid epidemic^16,29–31^. Historically, early studies suggested that the severe side effects of opioids, most notably respiratory depression, were primarily mediated by the *β*-arrestin signaling pathway, driving more than a decade of research aimed at discovering G-protein-biased mOR agonists^7,32^. This effort led to the development of supposedly biased agonists like *oliceridine* (TRV130) and *PZM21*^33,34^. However, recent rigorous reassessments have profoundly challenged the *β*-arrestin hypothesis^35^. Multiple independent studies have now shown that life-threatening adverse effects, such as respiratory depression, persist even in the absence of *β*-arrestin, and are likely mediated by the same G-protein-dependent pathways that provide analgesia^36^. Consequently, PZM21 and oliceridine have been shown to still induce respiratory depression and other typical opioid side effects. It is now increasingly accepted that the favorable therapeutic profiles of these drugs are primarily due to their low intrinsic efficacy (partial agonism) rather than true functional selectivity^35,37^. Specifically, partial agonists like buprenorphine exhibit a inherent ’ceiling effect’ that limits the maximum activation of pathways responsible for severe adverse effects, such as respiratory depression. This property effectively broadens the therapeutic window^38^, offering a safer separation between beneficial analgesia and on-target toxicity compared to full unbiased agonists. Yet, understanding the exact thermodynamic mechanisms and conformational landscapes linking partial agonism to this restricted signaling propensity remains a critical topic of vigorous research.

Recent advances in cryo-electron microscopy have provided high-resolution structures of MOR in active and inactive states, as well as in complex with either G-protein or *β*-arrestin1^39–43^. These structural studies have revealed key rearrangements associated with receptor activation in the presence of effector proteins,^24^ thus complementing the findings of NMR and molecular dynamics studies^16^. Taken together, these structural and dynamical studies highlight the importance of the intracellular displacement of helix 8 as the primary factor in selective binding to *β*-arrestin1 rather than G-proteins. However, a full characterisation of how biased and partial agonists affect the conformational energy landscape of MOR differently to full, unbiased agonists is still lacking, particularly with regard to the effector binding site. A quantitative description of the free-energy landscape governing MOR activation may allow us to discern if and how different ligand reshape signaling outcomes.

In this respect, enhanced sampling molecular dynamics (MD) simulations provide a powerful framework to address this challenge by enabling direct estimation of free-energy differences associated with large-scale conformational rearrangements inaccessible to conventional MD simulations^44–47^. In this work, we employ our GPCR-optimised enhanced sampling simulations to characterize the activation free-energy landscape of MOR.^48^. We compare the ligandless (apo) receptor with MOR bound to the endogenous agonist endomorphin-1 and to buprenorphine. Endomorphin-1 represents a physiological mode of receptor activation and serves as a reference for the native MOR activation pathway^49^, whereas buprenorphine is a partial agonist widely used both as an analgesic and as a first-line therapy for opioid use disorder^50^. Buprenorphine has a pharmacological profile distinct from that of full agonists such as DAMGO and morphine, including a ceiling effect for respiratory depression and altered efficacy across downstream signaling pathways^16,51–53^. Interestingly, our free-energy landscapes reveal that endomorphin-1 stabilizes two distinct basins within the active conformational ensemble of MOR, corresponding to receptor states compatible with G-protein and *β*-arrestin1 engagement, respectively. In contrast, in the presence of buprenorphine, the *β*-arrestin–compatible conformational basin is no longer observed as a thermodynamically stable minimum, indicating a loss of population of this signaling-competent state.

By mapping signaling-competent conformations onto the computed free-energy landscapes, we provide a thermodynamic description of how ligand binding modulates MOR signaling propensity. Our results establish a quantitative framework that links ligand-dependent conformational ensembles to effector-specific receptor states and offer mechanistic insight into the molecular determinants of opioid receptor signaling, with implications for the rational design of safer opioid therapeutics.

## Methods

### Systems Preparation

The structure of endomorphin-1-bound MOR was obtained from the G-protein-coupled MOR crystal structure deposited under PDB ID: 8F7R^54^ by removing the G-protein. A disulfide bridge was introduced between residues C^3.25^ and C^45.50^ to properly model MOR’s tertiary structure. To date, no experimentally resolved MOR bound to the agonist buprenorphine is available. For this reason, we exploited the refined structure of buprenorphine-bound MOR published by Gomes et al.^55^. Once modelled, both structures were embedded in a POPC/CHL membrane (80:20) using CHARMM-GUI^56^. The receptor N- and C-termini were capped with acetyl and methyl-amino protecting groups, respectively. The systems were solvated using the TIP4PD water model at 0.15 M NaCl and simulated with the DES-Amber force field^57^ in GROMACS 2023^58^. The buprenorphine molecule was parametrized using the GAFF2 force field^59^, while for the peptide endomorphin-1 the DES-Amber force field parameters were used. To equilibrate the systems, a thermalization protocol with progressively reduced positional restraints on heavy atoms was applied to relax potential structural artifacts while preserving the overall organization of the protein, ligands, and membrane. Equilibration consisted of consecutive 1 ns NVT and 1 ns NPT simulations at increasing temperatures from 100 to 300 K in 50 K increments with corresponding decrease in positional restraints on both the protein and the lipids. The last equilibration step consisted in a unbiased simulation of 200 ns at 300 K to fully hydrate and relax the system. Covalent bonds involving hydrogen atoms were constrained using LINCS^60,61^, enabling an integration time step of 2 fs. Short-range Coulomb and van der Waals interactions were truncated at 1.0 nm, and long-range electrostatics were treated with the particle-mesh Ewald method^62^. The temperature was maintained at 300 K using the velocity-rescale (v-rescale) thermostat^63^, and the pressure was controlled at 1 bar using the semi-isotropic c-rescale barostat^64^.

### OneOPES MD Simulations

The OneOPES sampling scheme^65^, a replica-exchange extension of OPES Explore^66^, was applied to sample the activation pathway of two systems systems, i.e., endomorphin-1-bound and buprenorphine-bound MORs. In this scheme, for each simulation eight exchanging replicas are run in parallel, the first one being convergence-dedicated (replica **0**) and the other seven being exploratory (replicas **1-7**). In all replicas, the main CV is biased with OPES Explore, while the exploratory replicas have additional OPES Explore biases on auxiliary CVs. The exploratory replicas are also progressively heated with OPES Expanded to facilitate the overcoming of hidden degrees of freedom and aid barrier crossing^67^ (see Tab. S1).

To properly describe GPCR activation, both the primary and auxiliary CVs must capture the slow degrees of freedom associated with this process, encompassing large-scale rearrangements of transmembrane helices (macroswitches) as well as localized conformational changes of conserved motifs (microswitches)^68^. To describe the global progression along the activation pathway, we employed a Euclidean PATH (EPATH hereafter) CV, as in our previous work^69^. The EPATH is defined in a two-dimensional space spanned by the RMSD with respect to the inactive and active MOR reference structures (i.e., PDB IDs 9MQJ^70^ and 8K9K^71^, respectively), where each milestone *i* is represented by a point (*RMSD_inactive_*_,*i*_; *RMSD_active_*_,*i*_). For the sake of clarity, the shape of EPATH CV is slightly curved above the diagonal in the (*RMSD_inactive_*; *RMSD_active_*) space, consistent with the activation pathway observed in previous works on GPCR activation^48,69^, which allows the system to traverse an entropic basin populated by conformations that differ substantially from both inactive and active reference states.

Consistent with our previous works^48,69^, auxiliary distance CVs were defined for key microswitch motifs, i.e., *PIF*, *DRY*, *NPxxY*, and *YY* to describe their side chains rearrangement^40,68^. These distances were biased in the exploratory replicas. Additionally, in all exploratory replicas, a hydration CV was included to sample the changes in the water network in the cytoplasmic cavity (see Fig. S1). The inclusion of this CV was supported by previous studies that demonstrated that water coordination at a specific site can substantially lower the activation free-energy barrier^72–76^ and that diverse GPCRs, including MORs, exhibit conserved water networks for stabilization and activation^77–79^. The simulations were run with GROMACS 2023 patched with PLUMED 2.9.1^80,81^. The details about thermostat, barostat, electrostatics cut-offs and force field are the same as described in the **Systems preparation** section. To ensure reliable statistics, each OneOPES simulation was run in triplicates, up to 900 ns. The results are thus reported with the corresponding average and standard deviation. Finally, we chose to re-project the free-energy results onto a simplified EPATH CV with seven milestones, to be consistent with our previous work on *apo-MOR* and aid the comparison of the results^69^. When re-weighting, the first 200 ns are discarded from the trajectory to remove the starting part of the simulation where the bias is strongly out-of-equilibrium. The full description of the OneOPES parameters, CVs, and the original biased path are reported in the **Supplementary Information**.

### Ligand–receptor contact analysis across helix 8 conformational pools

To assess ligand–receptor interactions in the ligand-bound MOR simulations, the contacts between the given ligand and MOR were computed separately for distinct conformational pools corresponding to different orientations of H8 relative to the intracellular surface. These pools, *H8_Barr* and *H8_Gi*, were defined based on their similarity with respect to recently released *β*-arrestin1-bound MOR structure (PDB ID: 9WSX ^24^) and the torsional angle measured between the C*α*s of the amino acids T^7.29^, A^7.54^, L^7.56^, F^8.54^. For the sake of clarity, we exploited 9WSX to build two CVs i.e., *RMSD_Barr_*_−*bound*_, defined as the RMSD on 9WSX’s C*α*s, and *RMSD_Barr_*_−*bound*−*H*8_, measured only on H8’s C*α*s (both were aligned on TM1-7’s C*α*s). For each conformational pool, a contact was defined when any ligand atom was within a distance cutoff of 4.0 Å from any atom of a given receptor residue^82^. The frequency of contacts occurrence was quantified as the fraction of frames in which the contact is present over the total number of frames within a given pool. Residues exhibiting the largest differences in frequency of occurrence between pools were identified by calculating pairwise differences in contact frequencies and selecting those with the highest variance between states. Contact analysis was carried out with MDAnalysis^83,84^.

## Results

### Ligand-dependent free-energy landscapes of MOR activation

To investigate how ligand binding modulates MOR activation, we employed enhanced sampling molecular dynamics simulations using the OneOPES framework to map the free-energy landscapes of the receptor in different functional states. In the following, we compare the ligandless receptor (*apo-MOR* hereafter), with MOR bound to the endogenous ligand endomorphin-1 and the partial agonist buprenorphine (*endo-MOR* and *bpnp-MOR*, respectively) (see Fig. 1a and Fig. S2).

**Figure 1:**
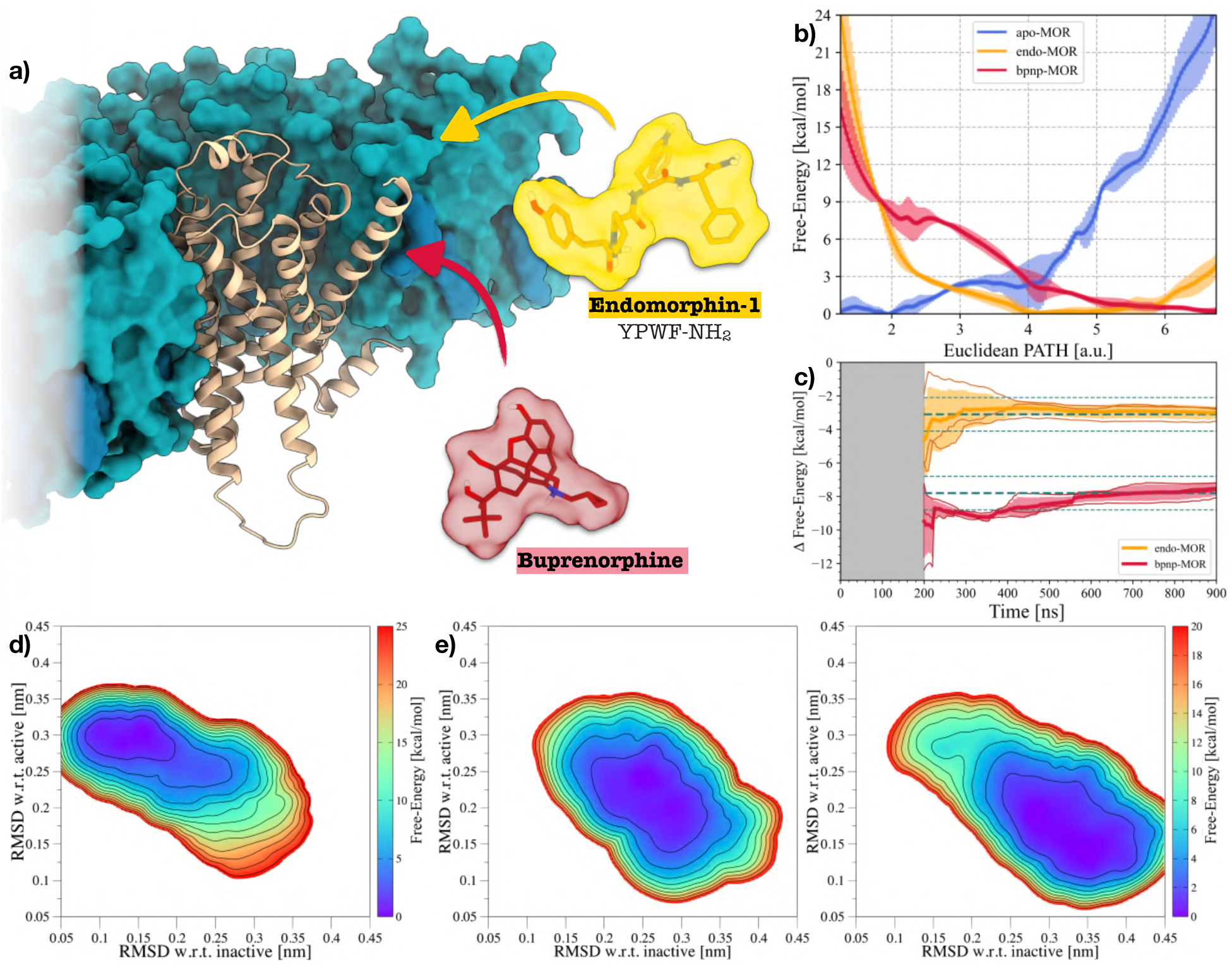
Effects of ligand-binding on MOR’s activation free-energy. **(a)** MOR (in tan) embedded into a POPC/CHL membrane and the two investigated agonists, i.e., endomorphin-1 (in orange) and buprenorphine (in red). POPC and CHL are represented via their solvent-exposed surface and colored in turquoise and teal, respectively. **(b)** 1D free-energy profile re-weighted along the EPATH CV. The profiles measured for *apo-MOR* ^48^, *endo-MOR*, and *bpnp-MOR* are displayed as solid lines and colored in blue, orange, and red, respectively. Standard deviations are shown as transparent areas and colored accordingly. **(c)** Evolution of the free-energy difference (Δ*G*) over simulation time. Each independent trajectory is shown as a solid line, whereas their average is represented as a thicker solid line. The transparent area indicates their standard deviation. **(d)** 2D FES reweighted in the (*RMSD_inactive_*; *RMSD_active_*) space for *apo-MOR*, coloured according to the color bar on the right side. **(e)** 2D FES reweighted in the (*RMSD_inactive_*; *RMSD_active_*) space for *endo-MOR* (right side), and *bpnp-MOR* (left side). Each FES was coloured according to the color bar on the right side.

The 1D free-energy profiles along the activation path, shown in Fig. 1b, reveal marked differences in the thermodynamic accessibility of active-like conformations across the three systems. In the apo receptor, the global minimum is located within the inactive region (EPATH ∼ 2), while active-like conformations form a higher-energy metastable state (Δ*G* ∼ 18 kcal/mol^48^). Endomorphin-1 significantly reshapes the free-energy landscape, leading to a stabilization of both the intermediate and active-like conformational pools, and making the active-like basin more accessible (Δ*G* ∼ −3 kcal/mol, see Fig. 1c). In contrast, buprenorphine produces a distinct free-energy landscape characterized by the full stabilization of both intermediate and activation-related conformations (Δ*G* ∼ −8 kcal/mol). Indeed, active-like states become more accessible compared to both *apo-MOR* and *endo-MOR*, and the global minimum is shifted towards intermediate-to active conformations (EPATH *>* 5), consistent with the partial agonist profile of buprenorphine.

To further characterize the structural features underlying these free-energy profiles, we re-projected the accumulated bias potentials on the (*RMSD_inactive_*; *RMSD_active_*) two-dimensional space, defined by structural similarity to inactive and active reference conformations (see Fig. 1d-f). This representation confirms the trends observed in the 1D EPATH free-energy profiles. *Apo-MOR* predominantly prefers conformations close to the inactive reference. *Endo-MOR*, instead, populates intermediate regions between inactive and active conformations. Finally, *bpnp-MOR* exhibits a broader conformational distribution that extends toward active-like states.

Together, the latter results show that the binding of these two ligands transforms the thermodynamic landscape of MOR activation and highlight distinct ligand-specific effects, that is, endomorphin-1 promotes activation by stabilizing intermediate structures along the activation pathway, whereas buprenorphine favors both intermediate and active conformational pools.

### Ligand-dependent effectors recruitment

The free-energy landscapes described in the previous section provide a global thermodynamic view of MOR activation and its ligand-dependent modulation. Nevertheless, to shed light upon the molecular determinants underlying these differences, we had to assess the behavior of the canonical GPCR microswitches involved in class-A GPCR activation. For this goal, we reweighted the accumulated bias potentials onto the CVs describing the microswitches and hydration dynamics as a function of the EPATH CV (see Figs. S3 and S4).

While the distances involved with the *PIF* and *DRY* motifs follow a similar pattern between *endo-MOR* and *bpnp-MOR*, differences arise in the behavior of the intracellular cavity. In detail, *endo-MOR* simulations exhibited a limited extent of motion for the Y^5.58^-Y^7.53^, ranging from ∼1.0 nm at EPATH < 2 to ∼0.8 nm at EPATH > 5. Conversely, *bpnp-MOR* presented a wider mobility: in the inactive state, the *YY* motif presents a clear minimum at ∼1.4 nm while, in the active state, two iso-energetic minima are stabilized by Y^5.58^-Y^7.53^ distances of ∼0.8 nm and ∼1.2 nm; a mechanism more akin to what we measured for *apo-MOR*^48^. This aspect reflects on the hydration of the binding cavity. In *endo-MOR*, the differences in hydration are almost iso-energetic across the activation landscape, indicating that the peptide-binding process facilitates the co-existence of different structural ensembles. In contrast, *bpnp-MOR* exhibits a well-defined hydration profile, characterized by transient water influx in pre-active conformations followed by partial dehydration upon stabilization of active-like states, as MOR points towards a well specific conformational state.

While microswitches capture key features of MOR activation, they do not directly resolve effector-specific conformational selection, and therefore cannot distinguish between receptor states competent for G-protein or *β*-arrestin engagement. To overcome this limitation, we next focus on large-scale allosteric rearrangements associated with effector-selective signaling, with particular emphasis on the intracellular orientation of helix 8 (H8). The recent cryo-EM structure of the MOR–*β*-arrestin1 complex (PDB ID: 9WSX) ^24^ provides a structural reference for a *β*-arrestin1–compatible state, in which H8 adopts a distinct orientation compared to the Gi-bound active conformation used as reference in this work (see Fig. 2a and **Methods** for further details). Building on this structural framework, we reprojected the accumulated OneOPES bias potentials for *endo-MOR* and *bpnp-MOR* onto a two-dimensional space defined by *RMSD_Barr_*_−*bound*_ and *RMSD_active_*, where *RMSD_Barr_*_−*bound*_ is computed with respect to the *β*-arrestin1-bound MOR structure (see Fig. 2b-c). For *endo-MOR*, this new analysis allows us to discriminate two different conformational pools sharing the same *RMSD_active_* values, i.e., *H8_Barr* and *H8_Gi*, with *H8_Barr* (the deeper minimum) more similar to PDB ID: 9WSX. Conversely, *bpnp-MOR* sampled a broader and more overlapping conformational ensemble, hinting a weaker energetic separation between the two H8 states. To better quantify the differences between *H8_Barr* and *H8_Gi*, we exploited the torsional angle between TM7 and H8 as a discriminant (see Fig. 2d and Fig. S5). The reweighted distributions (see Fig. 2e) show that *endo-MOR* populates two well-separated H8 orientations, while *bpnp-MOR* exhibits a distribution with substantial overlap between the *H8_Gi* and *H8_Barr* conformational pools. Representative structures extracted from these distributions (see Fig. 2f) better illustrate that the *H8_Gi*-like state corresponds to downward displacement of H8, away from the inner leaflet of the plasma membrane. These results suggest that MOR’s conformations more prone to binding *β*-arrestin1 are spontaneously sampled in a ligand-dependent manner, with different orthosteric ligands allosterically decreasing hidden energy barriers associated with the displacement of H8 and, consequently, effector recruitment.

**Figure 2:**
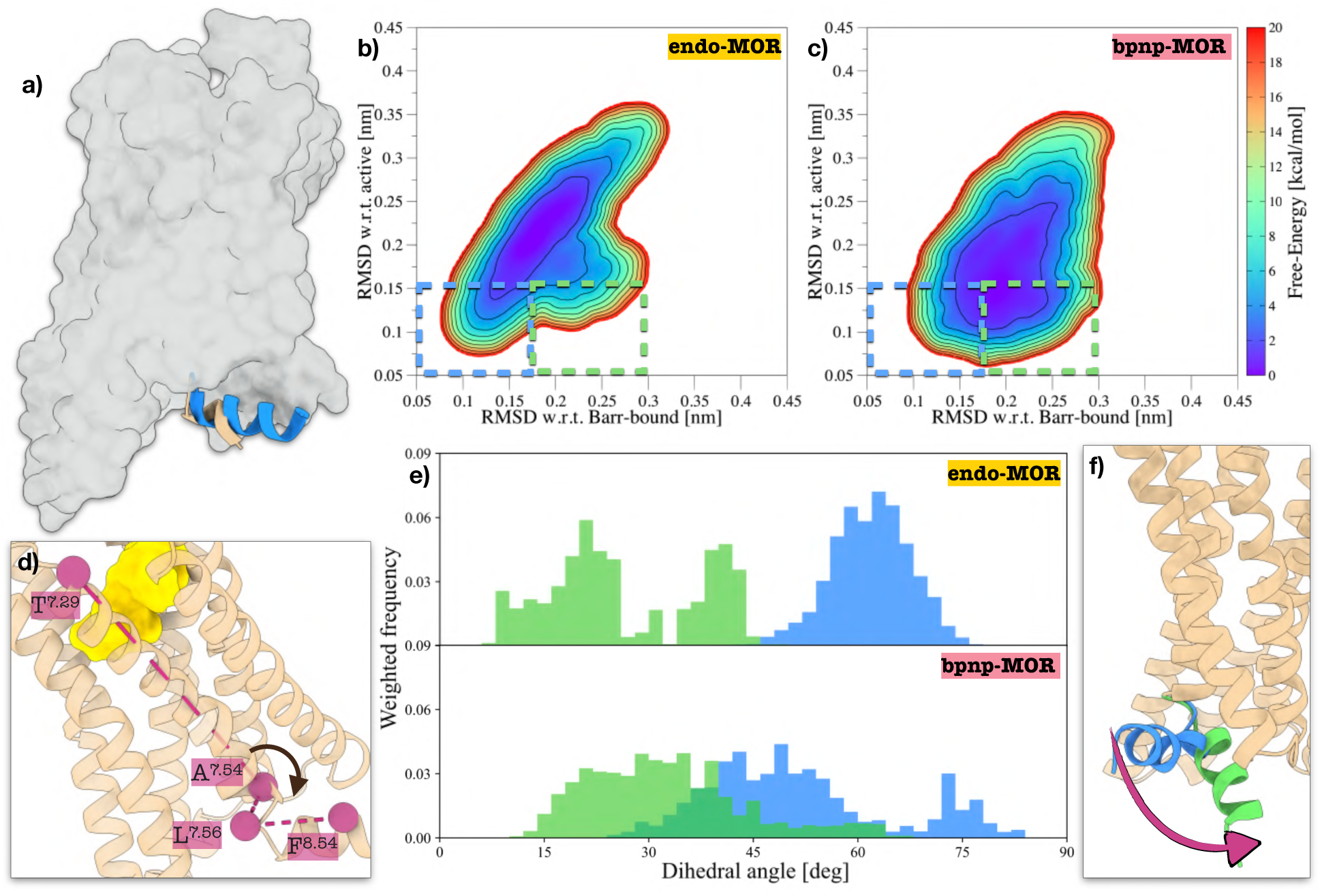
Exploration of β-arrestin-1 bound conformations of MOR. **(a)** Superimposition of the Gi-bound MOR structure used to built the EPATH CV (PDB ID: 8K9K, in tan) with the recently released *Barr*-bound MOR pdb structure (PDB ID: 9WSX ^24^, in blue). The TM1-TM7 bundle in common is represented as a surface colored in grey, while H8s are displayed as cartoon helices. **(b-c)** 2D FES reweighted in the (*RMSD_Barr_*_−*bound*_; *RMSD_active_*) space for *endo-MOR* and *bpnp-MOR*. The regions indicated by dashed lines show two different active-like conformational pools relative to H8 orientation, i.e., *H8_Barr* in blue and *H8_Gi* in green. **(d)** Schematic depiction of the torsional angle employed to measure the inclination of H8 with respect to the membrane. The selected atoms are showed as rigid spheres colored in magenta. **(e)** Re-weighted histograms of the torsional angle distributions in the conformational pools of *H8_Barr* (in blue) and *H8_Gi* (in green), for both *endo-MOR* and *bpnp-MOR*. **(f)** Representative structures of MOR’s H8 as extracted from the reweighted distributions showed in (**e**).

### Ligand’s position and interactions influences H8’s conformational pool

As hinted in the latter analysis, distinct MOR’s active-like conformations seem to arise from the coupling between the orthosteric binding site and H8. Thus, we analyzed ligand–receptor interactions patterns across the two active-like ensembles (*H8_Gi* and *H8_Barr*, as defined in Fig. 2b) to investigate if specific binding modes control this behavior. For *endo-MOR*, we observe a clear dependence of residue contact frequencies on the H8 conformational state (Fig. 3a). Residues located on TM3/TM5 display the largest variability, with I^3.40^, K^5.39^ and A^5.46^ emerging as key discriminants between the two conformational pools. The *H8_Barr* ensemble, corresponding to conformations in which H8 lies closer to the membrane, is characterized by frequent interactions between endomorphin-1’s Y1 and I^3.40^ and A^5.46^. In contrast, the *H8_Gi* ensemble features a displacement of Y1 towards the extracellular environment, with more frequent interactions with K^5.39^. In contrast, buprenorphine displays a markedly reduced variability in binding pose across *H8_Barr* and *H8_Gi* conformational ensembles, as evidenced by the nearly invariant contact patterns observed in both states (Fig. S6). To further characterize ligand-dependent differences during MOR activation, free-energy landscapes were projected onto the (*EPATH*; *RMSD_Barr_*_−*bound*−*H*8_) space (Fig. 3c-d), to better focus on H8’s . For *endo-MOR*, the landscape exhibits two distinct low-energy regions at EPATH > 6, with the deepest basin corresponding to H8’s conformations comparable to the *β*-arrestin1-bound structure (*RMSD_Barr_*_−*bound*−*H*8_ < 0.1 nm). In contrast, *bpnp-MOR* displays a broader and more continuous free-energy basin in the active-like portion of the EPATH CV, disfavouring *β*-arrestin1-bound structures. These results indicate that the two ligands differentially modulate the distribution of active-state conformations and the relative populations of intracellular signaling-competent states.

**Figure 3:**
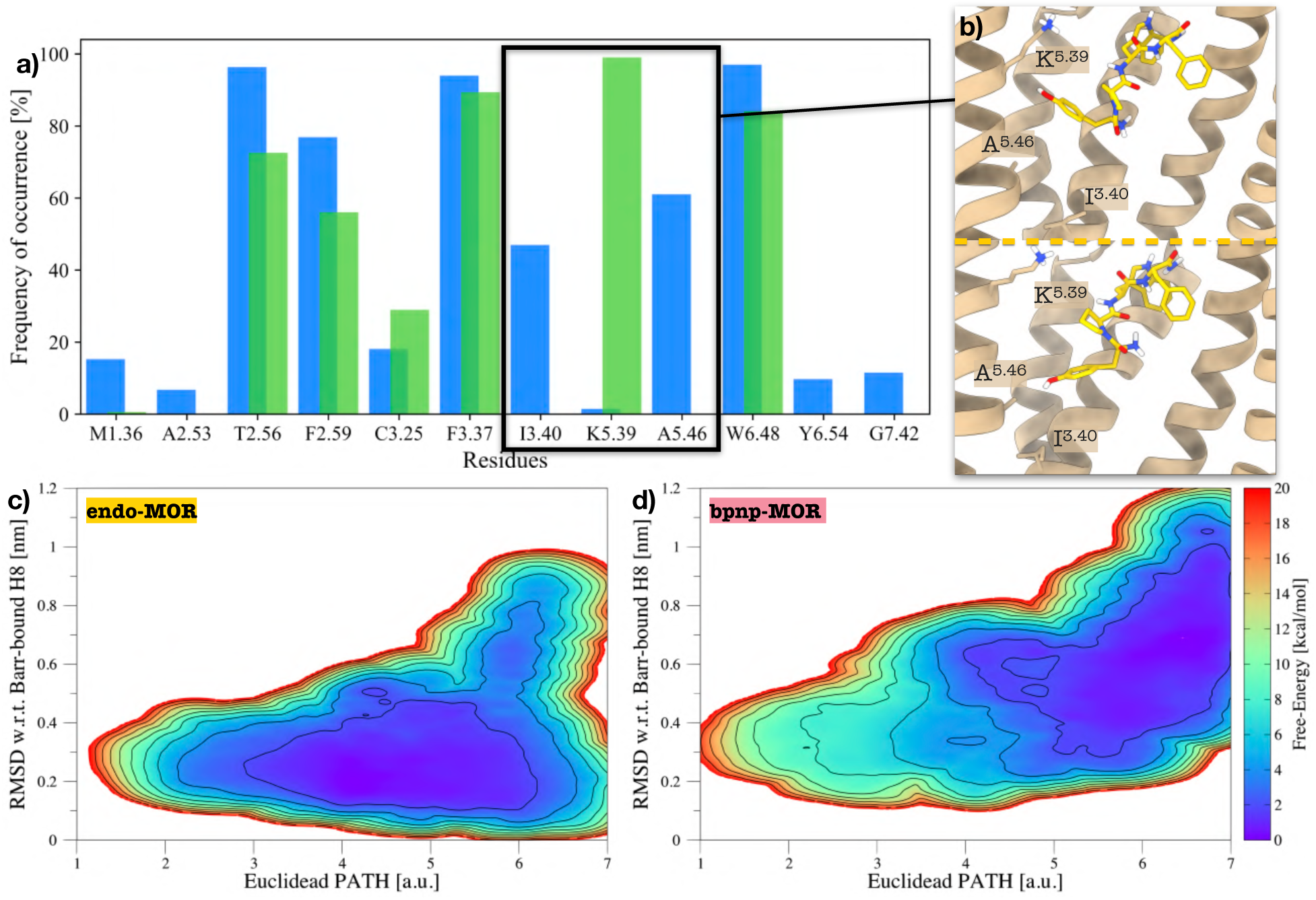
Analysis of ligands’ binding mode in MOR’s orthosteric binding site. **(a)** Frequency of occurrence of MOR’s most relevant residues binding endomorphin-1. The colors of the histogram bars show different active-like conformational pools relative to H8 orientation, i.e., *H8_Barr* (in blue) and *H8_Gi* (in green). The black square highlights I^3.40^, K^5.39^ and A^5.46^, the amino acids displaying the highest variability among H8’s conformational pools. **(b)** Representative binding poses of endomorphin-1 for *H8_Gi* (upper panel) and *H8_Barr* (lower panel). Endomorphin-1 is shown in orange, while MOR’s residues are coloured in tan. **(c,d)** 2D FES in the (*EPATH*; *RMSD_Barr_*_−*bound*−*H*8_) space for *endo-MOR* and *bpnp-MOR*.

## Discussion

Understanding how ligands encode specific signaling outcomes in GPCRs remains a central challenge in molecular pharmacology. In the case of the *µ*-opioid receptor, this question is particularly pressing, as subtle differences in signaling profiles underlie both therapeutic efficacy and severe adverse effects^85,86^. Despite the growing structural characterization of MOR in complexes with different effectors^24,39–42^, a quantitative framework linking binding of the ligand to thermodynamic accessibility of signaling-competent states remains elusive so far. In the present manuscript, we provide such a framework by combining OneOPES enhanced sampling simulations with a systematic mapping of conformational ensembles associated with distinct intracellular partners, i.e., G-protein and *β*-arrestin1. Moreover, we show that ligand binding reshapes the free-energy landscape by differentially modulating the accessibility of multiple active conformations. Notably, endomorphin-1 stabilizes two distinct active minima associated with receptor states compatible with G-protein and *β*-arrestin1 engagement, respectively, whereas buprenorphine selectively destabilizes the conformational pool prone to *β*-arrestin–binding, effectively restricting the accessible signaling ensemble. Another central finding of our study is that ligand-dependent differences in binding pose directly couple to the conformational variability of intracellular regions, in particular H8. For the endogenous agonist endomorphin-1, we observe a flexible binding mode that dynamically explores multiple geometries within the orthosteric pocket. This positional heterogeneity translates into a diversified set of local interactions, especially along TM5.

Within our framework and supported by recent literature^15,16,24^, H8 emerges as a hallmark of ligand-dependent allosteric effects. Notably, conformations in which H8 remains closely associated with the membrane are more consistent with receptor states competent for *β*-arrestin coupling. While we do not explicitly model effector binding, our results provide a thermodynamic basis for how ligand-dependent modulation of H8 accessibility could bias downstream signaling pathways. Future experimental and computational investigations will be needed to validate this mechanism and to clarify how TM5-mediated conformational changes contribute to the selective stabilization of effector-specific receptor states.

These insights are particularly timely in light of the ongoing opioid crisis, driven in large part by the widespread use of highly potent synthetic opioids such as fentanyl^87,88^.

Our results suggest that ligands can be designed to selectively reshape the conformational landscape of MOR, thereby favoring specific intracellular coupling modes. Such an approach may enable the development of safer opioid therapeutics by promoting beneficial signaling pathways while minimizing those associated with adverse effects^89^. More broadly, the framework introduced here is not limited to MOR but is readily extendable to other class-A GPCR, whose biased signaling properties are already exploited for therapeutic perspectives^90,91^. Indeed, by explicitly linking ligand-dependent free-energy landscapes to effector-specific conformational states, this approach can provide a general strategy to rationalize and predict signaling outcomes from first principles. We anticipate that integrating such thermodynamic descriptions with experimental measurements of signaling bias will be essential for advancing the rational design of next-generation GPCR-targeting drugs.

## Acknowledgment

This work was financially supported by the Swiss National Science Foundation and Bridge funding schemes (project numbers: 200021_204795, CRSII5_216587, and 40B2-0_203628). We acknowledge the Swiss National Supercomputing Centre for supercomputer time allocations on Daint (project ID: lp84 and lp166). We would like to thank Anton Ariel Hanke for useful discussion.

