## Supplementary Material for "Buprenorphine Restricts the Conformational Landscape of the *µ*-Opioid Receptor"

The data hereby reported are divided into the following sections:

- Supplementary Data 1: Details of the CVs employed for the OneOPES simulations;
- Supplementary Data 2: Sampling quality of the OneOPES simulations;
- Supplementary Data 3: Microswitches' behavior of MOR during the GPCR activation;
- Supplementary Data 4: Dependence on the TM7-H8 torsional angle for *endo-MOR* and *bpnp-MOR*;
- Supplementary Data 5: Binding modes of endomorphin-1 and buprenorphine in active states

### Supplementary Data 1

To investigate the activation pathway of the MOR GPCR in both the *endo-MOR* and *bpnp-MOR* systems, we employed a set of dedicated CVs reported in Tab. S1 and already employed in Ref. [1]. These variables are specifically designed to effectively capture both structural and hydration-related changes that occur during MOR activation, highlighting the complexity and significance of this vital biological process. As in Ref [2], the CVs on the residue couples (i.e., **D1-D7**) were built following a coarse-grained-inspired approach, treating the entire side-chain of specific amino acids as a putative dummy atom. For a detailed atomistic description of the **D1-D7** CVs, please refer to Fig. S2. The barrier of OPES Explore on the main CV is 50 kJ/mol for *endo-MOR* and 70 kJ/mol for *bpnp-MOR*. All the OPES MultiCV share a barrier of 3 kJ/mol.

| Replicas | 0 | 1 | 2 | 3 | 4 | 5 | 6 | 7 |
|---|---|---|---|---|---|---|---|---|
| OPES Explore | Neop1.s | Neop1.s | Neop1.s | Neop1.s | Neop1.s | Neop1.s | Neop1.s | Neop1.s |
| OPES MultiCV 1 | - | **D1**,yywo | **D1**,yywo | **D1**,yywo | **D1**,yywo | **D1**,yywo | **D1**,yywo | **D1**,yywo |
| OPES MultiCV 2 | - | - | **D2**,yywo | **D2**,yywo | **D2**,yywo | **D2**,yywo | **D2**,yywo | **D2**,yywo |
| OPES MultiCV 3 | - | - | - | **D3**,yywo | **D3**,yywo | **D3**,yywo | **D3**,yywo | **D3**,yywo |
| OPES MultiCV 4 | - | - | - | - | **D4**,yywo | **D4**,yywo | **D4**,yywo | **D4**,yywo |
| OPES MultiCV 5 | - | - | - | - | - | **D5**,yywo | **D5**,yywo | **D5**,yywo |
| OPES MultiCV 6 | - | - | - | - | - | - | **D6**,yywo | **D6**,yywo |
| OPES MultiCV 7 | - | - | - | - | - | - | - | **D7**,yywo |
| OPES MultiT | - | 301 K | 303 K | 306 K | 310 K | 317 K | 325 K | 335 K |

Table S1: The table presents the various CVs and parameters used in the OneOPES simulations. In the rows labelled "OPES Explore" and "OPES MultiCV", we outline the arrangement of the CVs across the replicas. The row titled "OPES MultiT" indicates the highest temperature reached during the trajectory, beginning from the thermostat's starting temperature of 300 K.

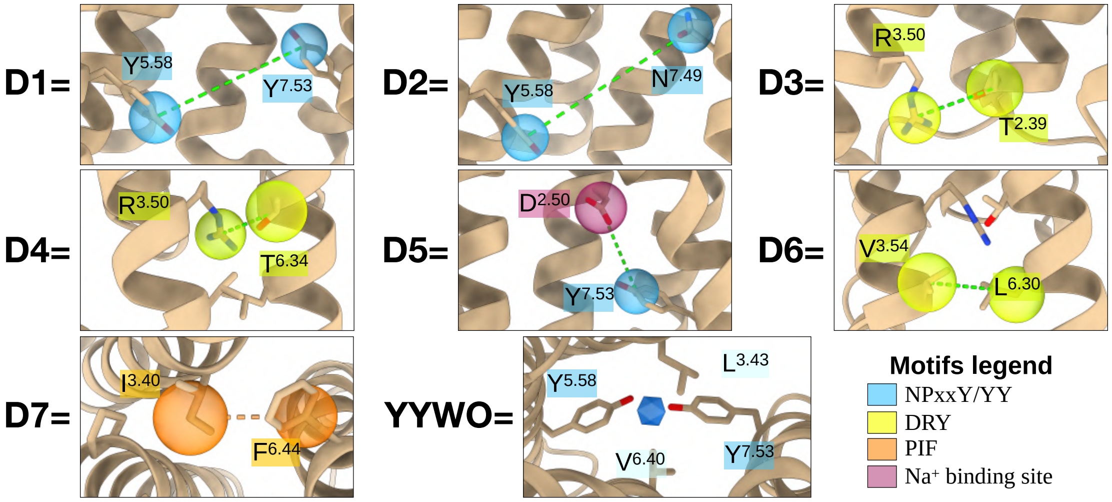


Figure S1: **Atomistic details of the CVs employed in the *apo-MOR*, *endo-MOR*, and *bpnp-MOR* OneOPES simulations.** For **D1-D7**, each CV is built as a distance between dummy atoms centered on the side-chains. The dummy atom built between $L^{3.43}$, $V^{6.40}$, and $Y^{7.53}$ and coordinating water molecules is represented as a blue icosahedron.

### Supplementary Data 2

In this section, we provide additional data on the behavior of the EPATH CV and the original EPATH* CV in the context of the *endo-MOR* and *bpnp-MOR* OneOPES simulations. For this purpose, we report Fig. S2, which illustrates the position of the milestones $M_i$ and $M_i^*$ within the $(RMSD_{inactive}; RMSD_{active})$ space and the 1D FES profile measured upon the EPATH* CV. Lastly, we present the sampling of the EPATH* CV from the *endo-MOR* and *bpnp-MOR* simulations, colored as a function of the accumulated bias potentials.

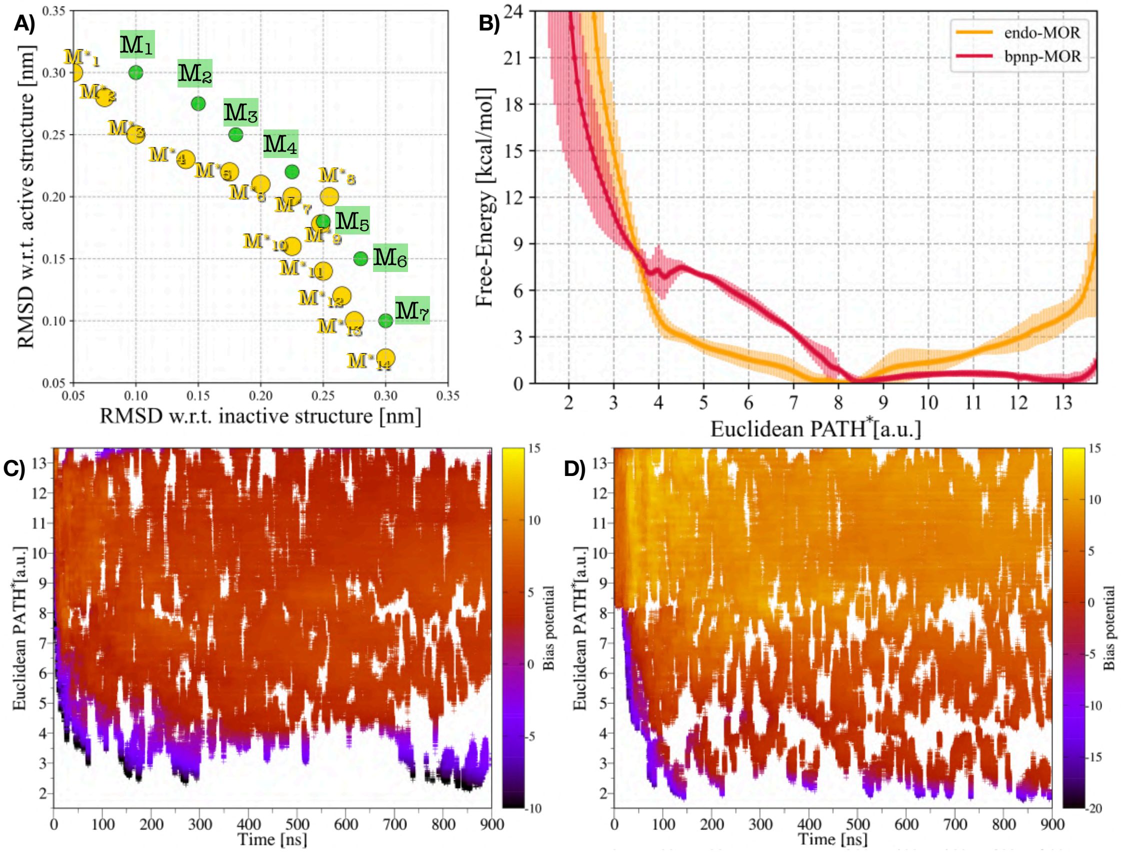


Figure S2: **Sampling of the main CVs in *endo-MOR* and *bpnp-MOR* OneOPES simulations. a)** Milestones $M_i^*$ sampled during the OneOPES simulations (i.e. the sampled EPATH*), and milestones $M_i$ of the EPATH CV upon which the outcomes coming from *endo-MOR* and *bpnp-MOR* have been re-weighted. **b)** Free-energy profile as a function of the EPATH* for *endo-MOR* and *bpnp-MOR*, averaged over three independent OneOPES simulations. The solid lines represents the mean free-energy, while the transparent shading indicates the standard deviation. **c-d)** Sampling of the EPATH* CV as a function of time in replica **0** for *endo-MOR* and *bpnp-MOR*, with data points colored according to the accumulated bias potential.

### Supplementary Data 3

In this section, we report a comparative analysis of MOR microswitch rearrangements obtained from the *endo-MOR* (left panel), and *bpnp-MOR* (right panel) OneOPES simulations. For each microswitch (i.e., the *NPxxY*, the *DRY*, and the *YY* motifs), we computed 2D FES as a function of the receptor's activation coordinate EPATH CV and the corresponding structural descriptor (see Fig. S3). At the same time, we also monitored the behavior of water molecules, whose accumulation in the intracellular cavity of the GPCR is a circumstance that favors the activation of the receptor (see Fig. S4). To provide reliable statistics, the reported 2D FES comes from the averages of the independent replicas.

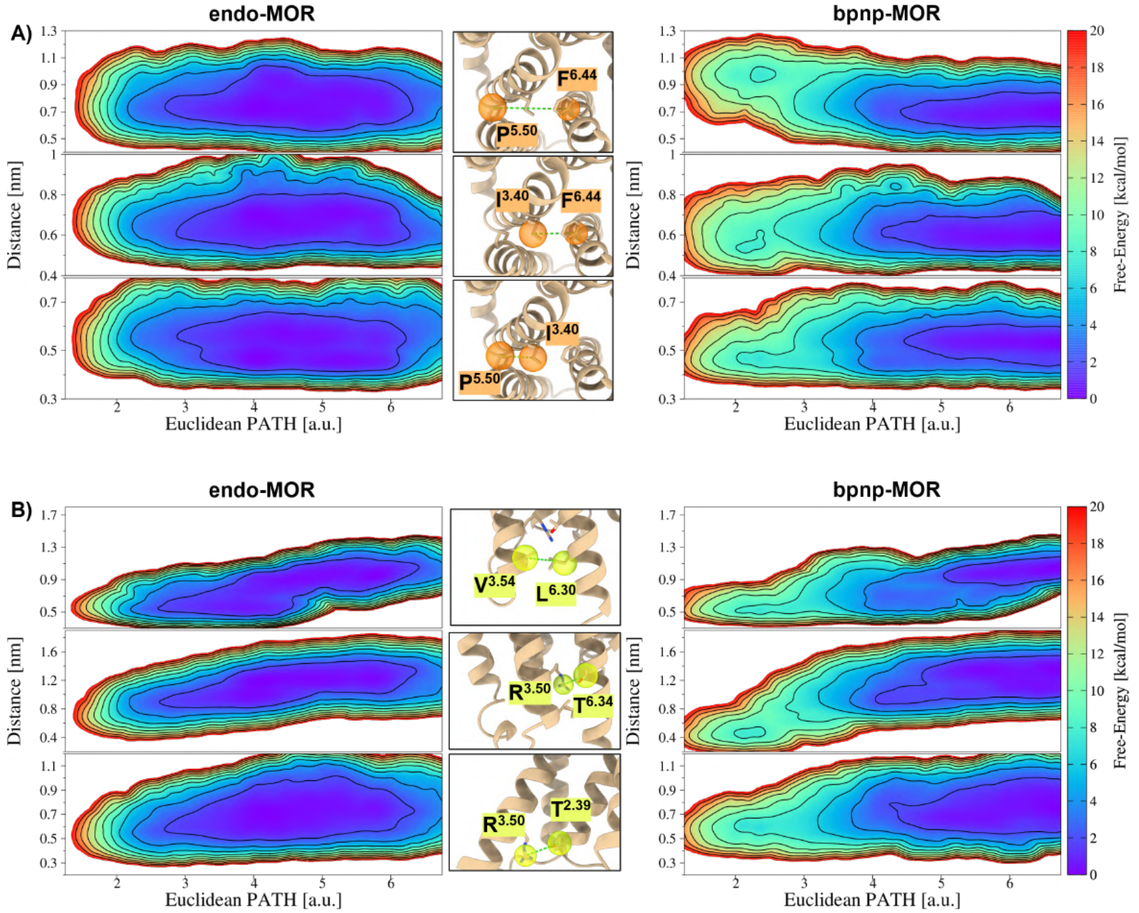


Figure S3: **Analyses of MOR's microswitches during the *endo-MOR* and *bpnp-MOR* OneOPES simulations. a)** 2D FES monitoring the Distance CVs for the residues of the *PIF* motif (i.e., $P^{5.50}$–$F^{6.44}$, $I^{3.40}$–$F^{6.44}$, and $P^{5.50}$–$I^{3.40}$) as a function of the EPATH CV for *endo-MOR* (left panel) and *bpnp-MOR* (right panel). **b)** 2D FES monitoring the Distance CVs for the residues in the proximity of the *DRY* motif (i.e., $V^{3.54}$–$L^{6.30}$, $R^{3.50}$–$T^{6.34}$, and $R^{3.50}$–$T^{2.39}$) as a function of the EPATH CV for *endo-MOR* (left panel) and *bpnp-MOR*(right panel). Each 2D FES has been coloured following the color scheme on the right side. Isolines are drawn every 2 kcal/mol.

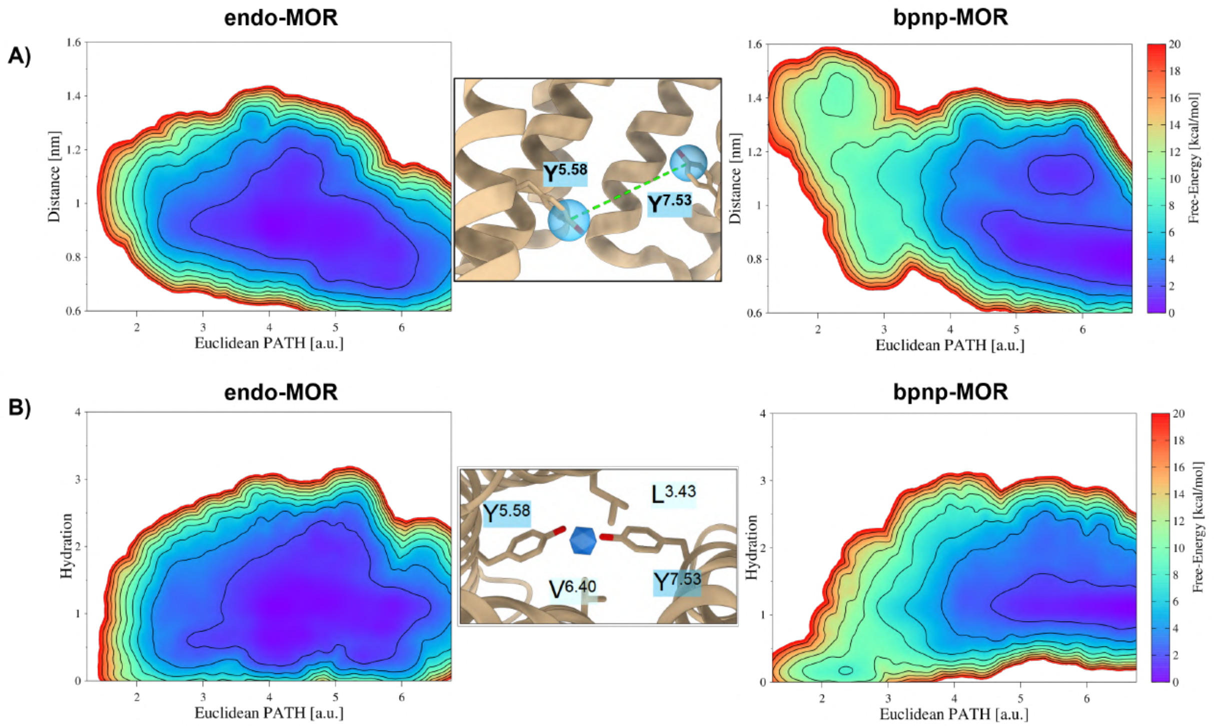


Figure S4: **Analysis of MOR's YY-motif and the hydration of the intracellular cavity during the *endo-MOR* and *bpnp-MOR* OneOPES simulations. a)** 2D FES monitoring the Distance CVs for the residues of the *YY* motif (i.e., $Y^{5.58}$–$Y^{7.53}$) as a function of the EPATH CV for *endo-MOR* (left panel) and *bpnp-MOR* (right panel). **b)** 2D FES associated with the hydration of MOR's intracellular cavity during the GPCR activation, as a function of the EPATH CV for *endo-MOR* (left panel) and *bpnp-MOR* (right panel). Each 2D FES has been coloured following the color scheme on the right side. Isolines are drawn every 2 kcal/mol.

#### Supplementary Data 4

In this section, we report the values of the torsional angle between TM7 and H8 mapped onto the ($RMSD_{Barr-bound}$; $RMSD_{active}$) 2D space. As discussed in the "Methods" section, the torsional angle is measured between the Cαs of the amino acids $T^{7.29}$, $A^{7.54}$, $L^{7.56}$, $F^{8.54}$ (see Fig. S5a). The other two panels show a representative replica for the *endo-MOR* and *bpnp-MOR* OneOPES simulations.

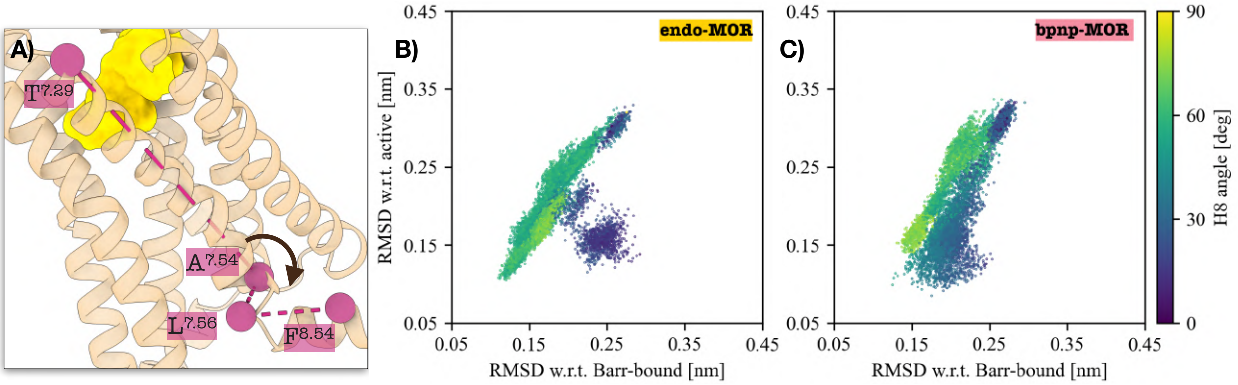


Figure S5: **Torsional angle between TM7 and H8 during the OneOPES simulations. a)** Definition of the torsional angle, as reported in Fig. 2. **b-c)** Re-projection onto the ($RMSD_{Barr-bound}$; $RMSD_{active}$) 2D space of the torsional angle for *endo-MOR* (**b**) and *bpnp-MOR* (**c**). Each point is coloured according to the colorgradient on the right side.

### Supplementary Data 5

In this section, we reported the complete interaction patterns established between MOR and the bound ligands within both active-state ensemble (i.e, *H8_Barr* and *H8_Gi*). Histograms reveal markedly different behaviors for endomorphin-1 and buprenorphine (see Fig. S5). Endomorphin-1 exhibits substantial variability in its interaction network across the different active-state conformations. As also discussed in the main text, this plasticity of the binding mode suggests a dynamic coupling between orthosteric-site interactions and the structural rearrangements associated H8's displacement (see Fig. S5a).

Conversely, buprenorphine maintains a largely conserved interaction pattern across all active-state ensembles. The majority of ligand–receptor contacts remain preserved despite the conformational differences observed within the receptor, indicating that buprenorphine occupies the orthosteric binding pocket in a relatively stable manner (see Fig. S5b). This behavior suggests a weaker dependence of the binding mode on the specific active-state conformation sampled by MOR.

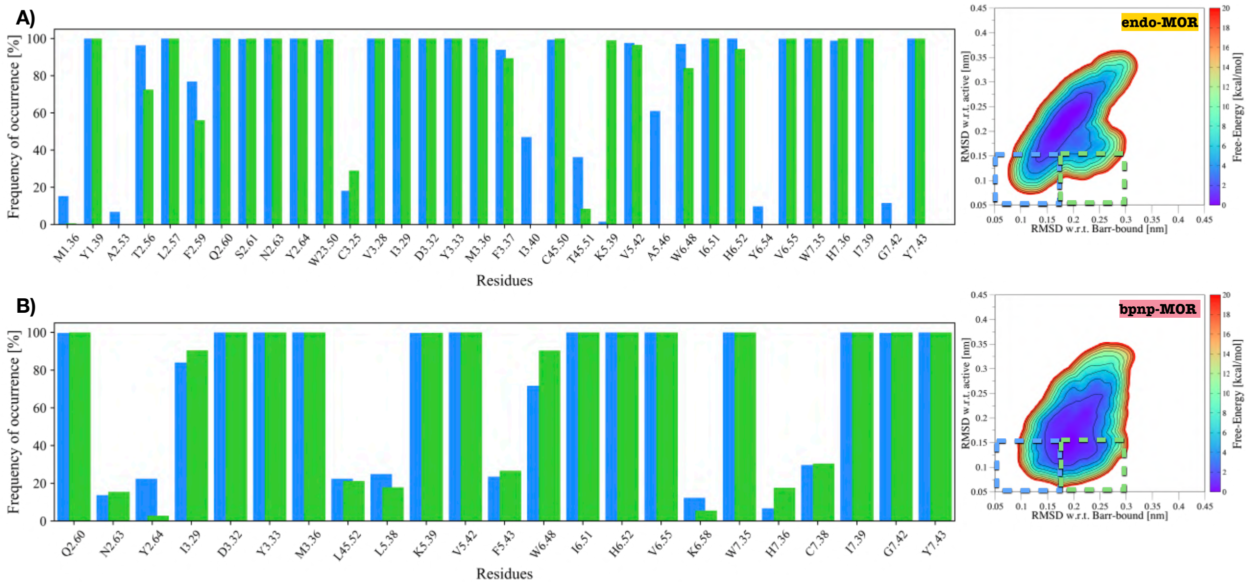


Figure S6: **Analysis of ligands' binding modes in MOR's orthosteric binding site for *endo-MOR* and *bpnp-MOR* OneOPES simulations. a-b)** Frequency of occurrence of MOR's amino acids binding endomorphin-1 (**a**) and buprenorphine (**b**). The colors of the histogram bars show different active-like conformational pools relative to H8 orientation, i.e., *H8_Barr* (in blue) and *H8_Gi* (in green), as in Fig. 3. On the right side, the 2D FES in the ($RMSD_{Barr-bound}$; $RMSD_{active}$) space upon which we defined the *H8_Barr* and *H8_Gi* conformational pools (as in Fig. 2).
